# Proteolytic program-dependent functions are impaired in INF2-mediated focal segmental glomerulosclerosis

**DOI:** 10.1101/530642

**Authors:** Balajikarthick Subramanian, Justin Chun, Chandra Perez, Paul Yan, Isaac Stillman, Henry Higgs, Seth L. Alper, Johannes Schlondorff, Martin R. Pollak

## Abstract

Regulation of the actin cytoskeleton is critical for normal glomerular podocyte structure and function. Altered regulation of the podocyte cytoskeleton can lead to proteinuria, reduced kidney filtration function and focal segmental glomerulosclerosis (FSGS). Mutations in inverted formin 2 (INF2), a member of the formin family of actin regulatory proteins, are the most common cause of autosomal dominant FSGS. INF2 is a multi-domain protein regulated by interaction between its N-terminal Diaphanous Inhibitory Domain (DID) and its C-terminal Diaphanous Auto-regulatory Domain (DAD). Although many aspects of the INF2 DID-DAD interaction are understood, it remains unclear why disease-causing mutations are restricted to the DID and how these mutations cause human disease. Here we report a proteolytic cleavage in INF2 that liberates the INF2 N-terminal DID to function independently of the INF2 C-terminal fragment containing the DAD domain. N-terminal DID region epitopes are differentially localized to podocyte foot process structures in normal glomeruli. This N-terminal fragment localization is lost in INF2-mediated FSGS, whereas INF2 C-terminal fragment epitopes localize to the podocyte cell body in both normal and disease conditions. INF2 cleavage is mediated by cathepsin proteases. In cultured podocytes, the wild-type INF2 N-terminal fragment localizes to membrane regions and promotes cell spreading, while these functions are impaired in a disease-associated INF2 mutant R218Q in the DID. These features are dependent on INF2-cleavage, with accompanying interaction of INF2 N-fragment with mDIA1. Our data suggest a unique cellular function of the DID dependent on INF2 cleavage and help explain the altered localization of FSGS-associated INF2 mutant polypeptides.

## INTRODUCTION

Glomerular epithelial cells, or podocytes, exhibit a polarized morphology characterized by a large cell body with extending primary processes and secondary foot processes (1, 2). These foot processes interdigitate to form uniquely specialized cell-cell contacts called slit diaphragms. The unique morphology of podocyte structure is dependent on the spatial and temporal regulation of the actin cytoskeleton which, if impaired, can cause proteinuric kidney diseases (3–5).

FSGS is etiologically and genetically heterogeneous (6). Highly penetrant Mendelian forms of FSGS are rare examples where we can unequivocally say we know the cause of disease (7). Although rare in absolute terms, INF2-associated FSGS is among the most common forms of inherited FSGS (8–10). Missense mutations in INF2 lead to kidney disease characterized by proteinuria, progressive kidney dysfunction, and FSGS with or without Charcot-Marie-Tooth disease (CMT) (11, 12). Approximately 9 to 17 percent of familial FSGS patients have INF2 mutations (7). INF2 is one of the 15 members of formin family of proteins, which share a formin homology domain (FH2) involved in control of actin polymerization (13, 14). However, most formins, including INF2, contain additional distinct regions of homology outside of the FH2 domain that are critical for actin assembly and interactions with regulatory binding partners. These regions include the diaphanous autoregulatory domain (DAD) near the INF2 C-terminus and the diaphanous inhibitory domain (DID) near the INF2 N-terminus. An intermolecular interaction between the DID and DAD maintains INF2 in an autoinhibited state (15). All pathogenic FSGS mutations of INF2 identified to date localize to the DID (16)

Cells express at least two isoforms of INF2, CAAX and non-CAAX (17, 18). The CAAX isoform localizes to ER-rich regions, whereas the non-CAAX form is cytoplasmic. Over the past few years, multiple studies have increased our understanding of INF2 isoform-mediated regulation of the actin cytoskeleton (15, 19–23). Many of these studies have relied on the cooperative regulation of the INF2 DID-DAD interaction to explain INF2 activity, as exemplified by INF2 regulation of mitochondrial fission (23–25). INF2 also modulates activity of other formins such as the mDIA subfamily, and promotes stable microtubule assembly (19, 26, 27). While disease-associated INF2 mutations appear to alter the DID-DAD interaction, disruption of this interaction does not explain why such mutations are limited to the DID. We reasoned that the DID region of INF2 might have unique functions in glomeruli independent of the DAD.

In the present study, we report the existence of proteolytic cleavage of INF2 that causes the DID to localize and function independently of the DAD and FH2 regions. Furthermore, the ability of wild-type INF2, but not mutant R218Q, to counteract mDIA activity and promote cell spreading in a cleavage-dependent manner indicates that INF2 possesses unique cleavage-dependent functions mediated via the N-terminal fragment. These activities are lost in the presence of pathogenic INF2 mutations.

## RESULTS

### INF2 is cleaved into two fragments separating the DID and DAD domains

We reasoned that if the INF2 DID possesses unique functions, these might be mediated via additional splice variants of INF2 that span just the N-terminus, as the annotated human genome (visualized using Ensembl or the UCSC Genome Browser) suggests the existence of such transcripts (28, 29). To examine this experimentally, we looked for the splice isoforms of INF2 present in human podocytes. mRNA expression analysis showed that both an INF2-CAAX isoform and a short isoform containing the first 5 exons of INF2 are expressed in human podocytes (Supplementary Figure 1A-C). Similar patterns of INF2 transcript expression were confirmed in human and mouse glomeruli (Supplementary Figure 1D). When we examined INF2 protein expression by immunoblot, we observed a band corresponding to full-length INF2 (~170 kDa) but no polypeptide band of the size encoded by the short transcript (expected at size ~ 25-35 kDa), either in cells or human glomerular lysate samples (blue box highlight in Supplementary Figure 1E).

However, using an antibody directed against the INF2 N-terminus, we observed an additional band of ~60 kDa in human glomerular lysates (Supplementary Figure 1E and Figure 1A). Using an antibody directed against the INF2 C-terminus, we also observed an additional band of approximately 100 kDa size (Figure 1A). Neither of these bands were present in INF2 CRISPR KO podocytes (Figure 1A). We saw similar INF2 immunoblot patterns in lysates from human glomeruli and primary human podocytes, as well as in glomeruli from wild-type mice, but not INF2 knockout mice (Figure 1A and B). When we overexpressed the INF2-CAAX isoform as GFP-INF2 or GFP-INF2-FLAG and probed for GFP or FLAG by immunoblot, we saw similar patterns (Figure 1C). Together, this data suggested specific cleavage in INF2 into two fragments, with the C-terminal fragment migrating at ~100 kDa and the N-terminal fragment migrating at ~60 kDa (~ 87kDa as the GFP-fusion) (Figure 1C).

**Figure 1.**
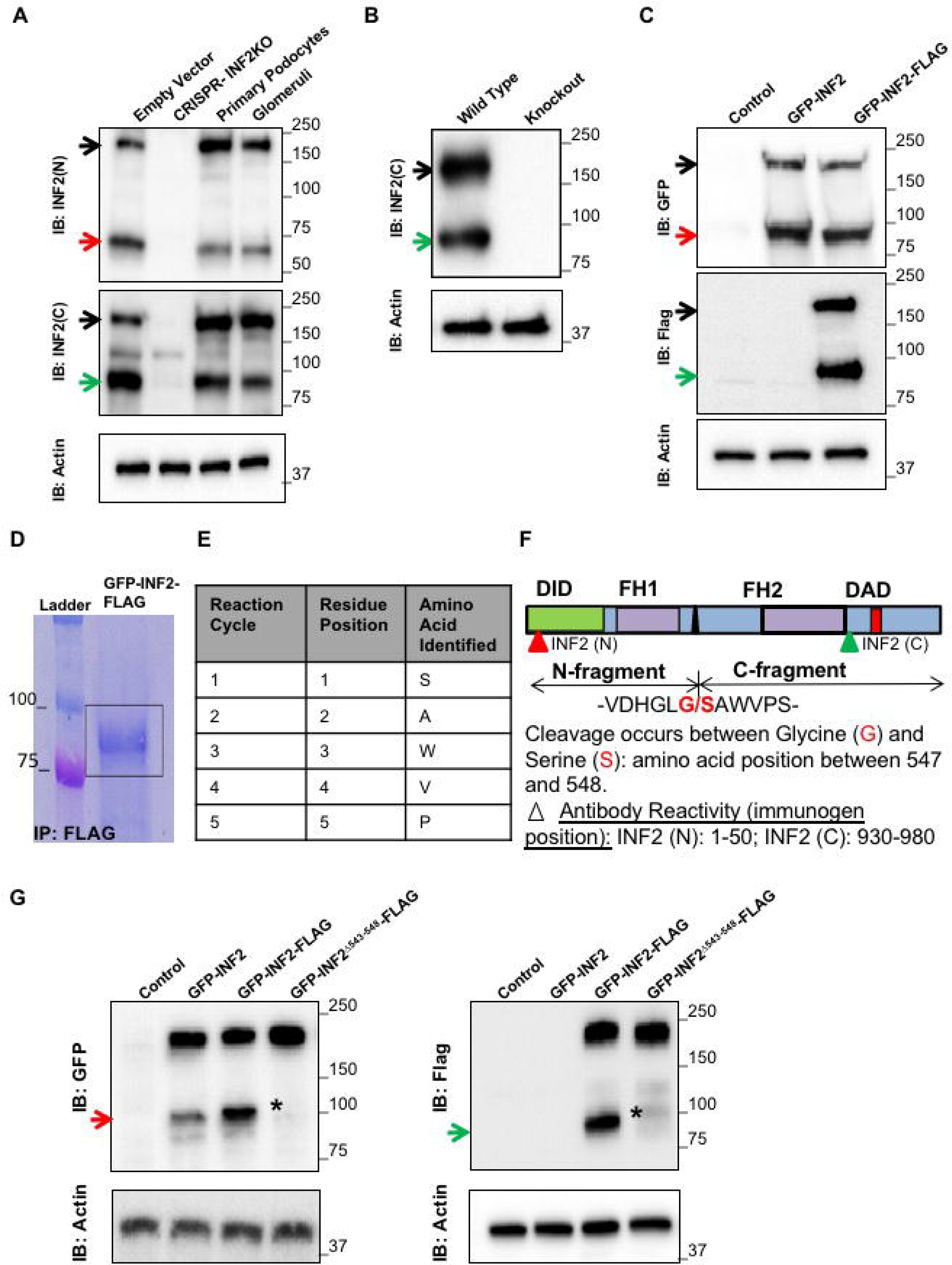
INF2 is cleaved into two fragments separating the DID and DAD regions. (A and B) Immunoblot analysis of INF2 in human and mouse samples. (A) Total cell lysates from primary mouse podocytes expressing empty vector (control), CRISPR INF2 KO mouse podocytes, primary human podocytes, and human glomerular tissue lysates were each probed with anti-INF2 antibodies specific for either the INF2 N-terminal (INF2(N)) or C-terminal-regions (INF2(C)). (B) Glomerular lysates from wild-type and *Inf2*-knockout mice were probed using a C-terminal-specific antibody. Arrows indicate different INF2 bands: Black - full-length INF2; Red - band detected only by N-terminal antibody; Green - band detected only by C-terminal antibody. Actin was used as a loading control. (C) Expression analysis of control vector, GFP-INF2, and GFP-INF2-FLAG constructs. Immunoblotting for GFP and FLAG tags detected similar full-length and smaller bands (red- GFP and green-FLAG), suggesting that INF2 is cleaved into two fragments. (D-F) Mapping of proteolytic cleavage site by N-terminal protein sequencing. (D) Coomassie stained PVDF membrane showing immunoprecipitated GFP-INF2-FLAG. Box indicates the immunoprecipitated C-fragment. (E) Amino acids obtained from first 5 cycles of N-terminal sequencing of the C-fragment. (Similar results were obtained from two independent immunoprecipitations and sequencing analyses). (F) Mapping of the INF2 cleavage site. The cleavage occurs between the amino acids highlighted in red, separating the DID and DAD regions, respectively, into N- and C-terminal fragments. (G) Validation of the cleavage site. Deletion of the cleavage site (Δ543-548) in GFP-INF2-FLAG results in complete loss of INF2 fragments. Red arrow, N-fragment; green arrow C-fragment. * indicates loss of cleaved fragments. Immunoblots shown are representative of three independent experiments.

To map the cleavage site, we overexpressed GFP-INF2-FLAG and immunoprecipitated the C-terminal fragment using an antibody to the FLAG tag, followed by N-terminal sequencing. The first five amino acids identified were “S-A-W-V-P” (Figure 1D). This sequence matched a unique locus on the C-terminal side of the INF2 DID (Figure 1E) and aligned with an *in silico*-predicted cathepsin K cleavage site spanning residues from 543-548. Cleavage was predicted to occur between INF2 residues glycine (G) 547 and serine (S) 548. To verify this cleavage site, we overexpressed a mutated form of GFP-INF2-FLAG in which we deleted the predicted cathepsin K cleavage site (non-cleavable GFP-INF2-FLAG). Cleavage fragments using this construct were no longer detectable by immunoblot (Figure 1F). This observation, along with the N-terminal sequencing results, confirmed that INF2 cleavage occurs between residues Glycine 547 and Serine 548.

### INF2 localization in glomeruli from normal and diseased kidney tissues correlates with proteolytic cleavage

To examine whether INF2 cleavage has an *in vivo* human correlation, we used structured illumination microscopy (SIM) to determine INF2 distribution within glomeruli in kidney biopsy samples from individuals without known kidney disease, as well as from individuals with INF2-mediated FSGS (Figure 2), Alport syndrome, and systemic lupus erythematosus (lupus nephritis) (Supplementary Figure 3). We stained these samples with both N-terminal-specific and C-terminal-specific INF2 antibodies to compare cleavage fragment localizations.

**Figure 2.**
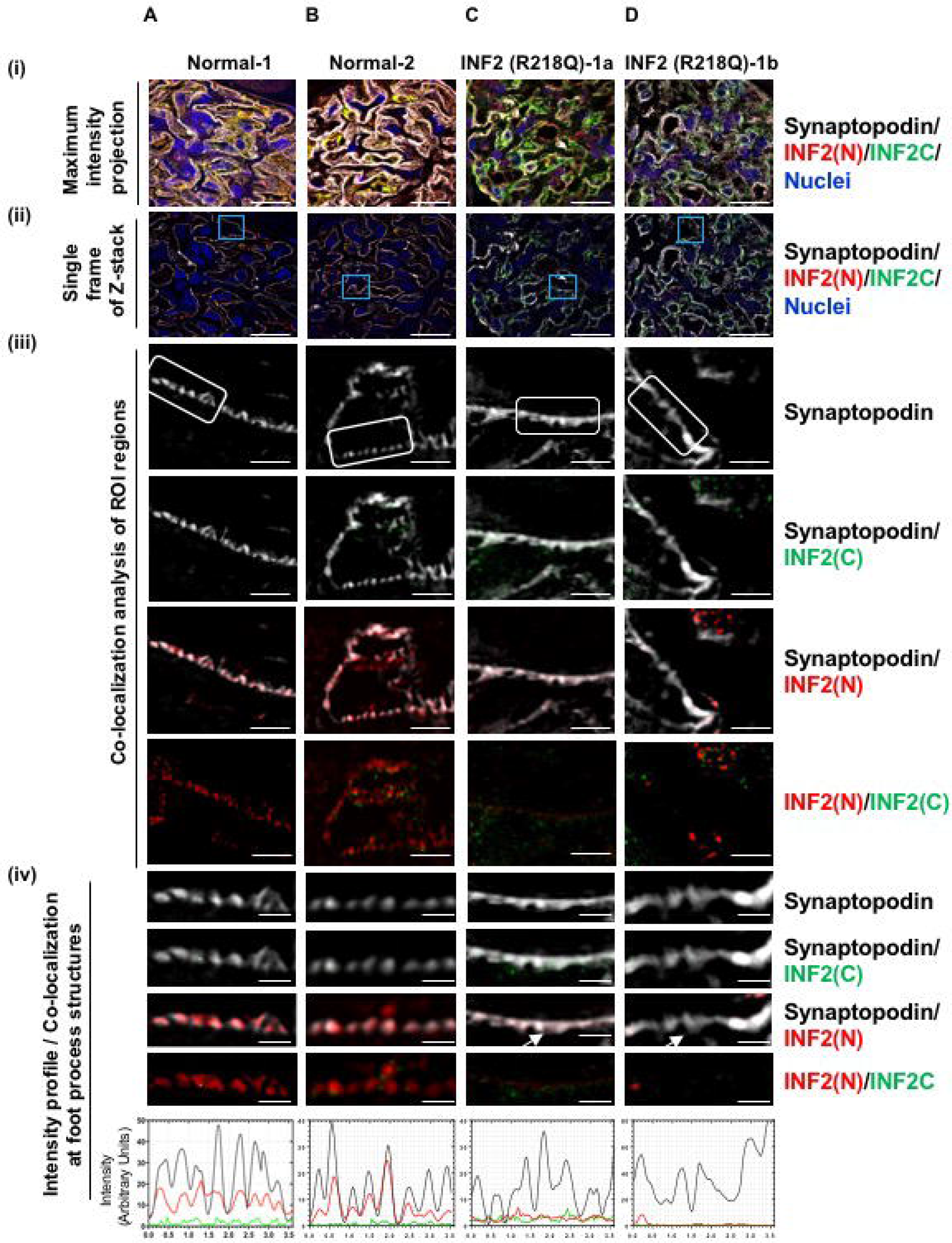
INF2 expression and localization in normal and diseased glomeruli by structured illumination microscopy (SIM). Kidney sections from normal individuals (A&B) and INF2 R218Q-associated FSGS (C&D) were stained for synaptopodin (grey), INF2 N-terminal region (N, red), INF2 C-terminal region (C, green), and nuclei (blue). Two representative glomeruli from two normal individuals and two diseased glomeruli from a single R218Q INF2-mediated FSGS patient are shown. (i & ii) SIM-processed low magnification micrographs. (i) Maximum intensity projection of 3D Z-stack optical frames. Scale bar 20 μm. (ii) A representative single optical frame of the 3D Z-stack shown in Scale bar 20 μm. (i). Blue boxes highlight Region of Interest (ROI) used for co-localization analysis. (iii) Co-localization analysis of INF2 with synaptopodin. Scale bar 2.5 μm. White boxes highlight foot process structures. INF2 N-terminal and INF2 C-terminal antibody staining patterns were differentially localized. (iv) Co-localization and corresponding intensity profiles of INF2 and synaptopodin in foot process structures. Scale bar 0.7 μm. INF2 N terminal staining co-localized with synaptopodin in normal foot process structures (A-iv and B-iv). This colocalization is lost in effaced regions of R218Q INF2 samples (arrow highlights; C-iv and D-iv). Intensity profiles confirm co-localization of fluorescent signal intensities.

Consistent with our previous observations in mice (30) and as shown in Supplementary Figure 3, we noted INF2 expression exclusively in glomerular podocytes of both normal and disease samples (Figure 2A-D). However, we observed discrete staining patterns for the different INF2 antibodies. In normal human kidney tissue, the staining pattern of C-terminal antibody showed predominant localization to the podocyte cell body, whereas the N-terminal antibody localized to both the podocyte cell body and foot process structures, co-localizing with synaptopodin (Figure 2A and B). By contrast, kidney tissue from an individual with FSGS due to INF2 mutant R218Q was remarkable for loss of N-terminal antibody staining from foot processes, whereas localization of C-terminal antibody staining in the cell body remained grossly unaltered (Figure 2C and D). Similar patterns were observed in the podocytes from individuals with Alport syndrome and with lupus nephritis (Supplementary Figure 2A and B). The segregation of N-terminal and C-terminal antibody staining patterns in normal kidney (Figure 2A and B), with localization of the N-terminal INF2 fragment to foot processes in normal samples and its loss in disease conditions, suggests that cleavage occurs *in vivo* and that loss of INF2 N-fragment-associated activity may lead to altered podocyte structure and function.

### INF2 cleavage is mediated by cathepsins and is not altered in the presence of FSGS-associated INF2 mutations

We reasoned that understanding the difference between INF2 N-terminal antibody staining in normal and diseased glomeruli might help elucidate the unique functions of the DID, as well as the mechanisms of INF2-associated FSGS. We hypothesized that the difference in N-terminal antibody staining of podocyte foot processes between normal and diseased kidney might reflect either (1) altered INF2 cleavage (e.g., mutations or other non-genetic factors affecting cleavage) or (2) events downstream of cleavage that affect N-fragment localization to foot processes. Therefore, we evaluated the regulation of INF2 cleavage. We first performed a chemical screen in podocytes using a protease inhibitor library to examine changes in the ratio of full-length INF2 to INF2 N-terminal fragment by immunoblot (Figure 3A). Most of the protease inhibitors tested had minimal effects on INF2 cleavage, whereas cathepsin inhibitors significantly increased the cleavage ratio, with greatest effect using cysteine protease inhibitor E64 (Figure 3B). To further confirm INF2 cleavage by cathepsin family members, we performed *in vitro* biochemical cleavage assays with cathepsin proteases and immunoprecipitates of wild-type full-length and non-cleavable mutant full-length forms of GFP-INF2 (Figure 3C). The immunoprecipitate of wild-type full-length INF2 included both full-length form and endogenously cleaved N-fragment of INF2, while the non-cleavable INF2 exhibited the full-length form alone (Figure 3C). The results from the biochemical assay showed that all tested cathepsins (B, L, and K) specifically cleaved the wild-type full-length GFP-INF2 yielding ~100 kDa C-terminal fragments and an increase in the levels of ~90 kDa N-terminal GFP-fusion fragments. INF2 proteolysis following cathepsin incubation was absent in the presence of cysteine protease inhibitor E64. By contrast, none of the tested cathepsins cleaved GFP-INF2 mutated to lack the cleavage site, confirming both the specificity of the cleavage site and the role of cathepsins in mediating this cleavage.

**Figure 3:**
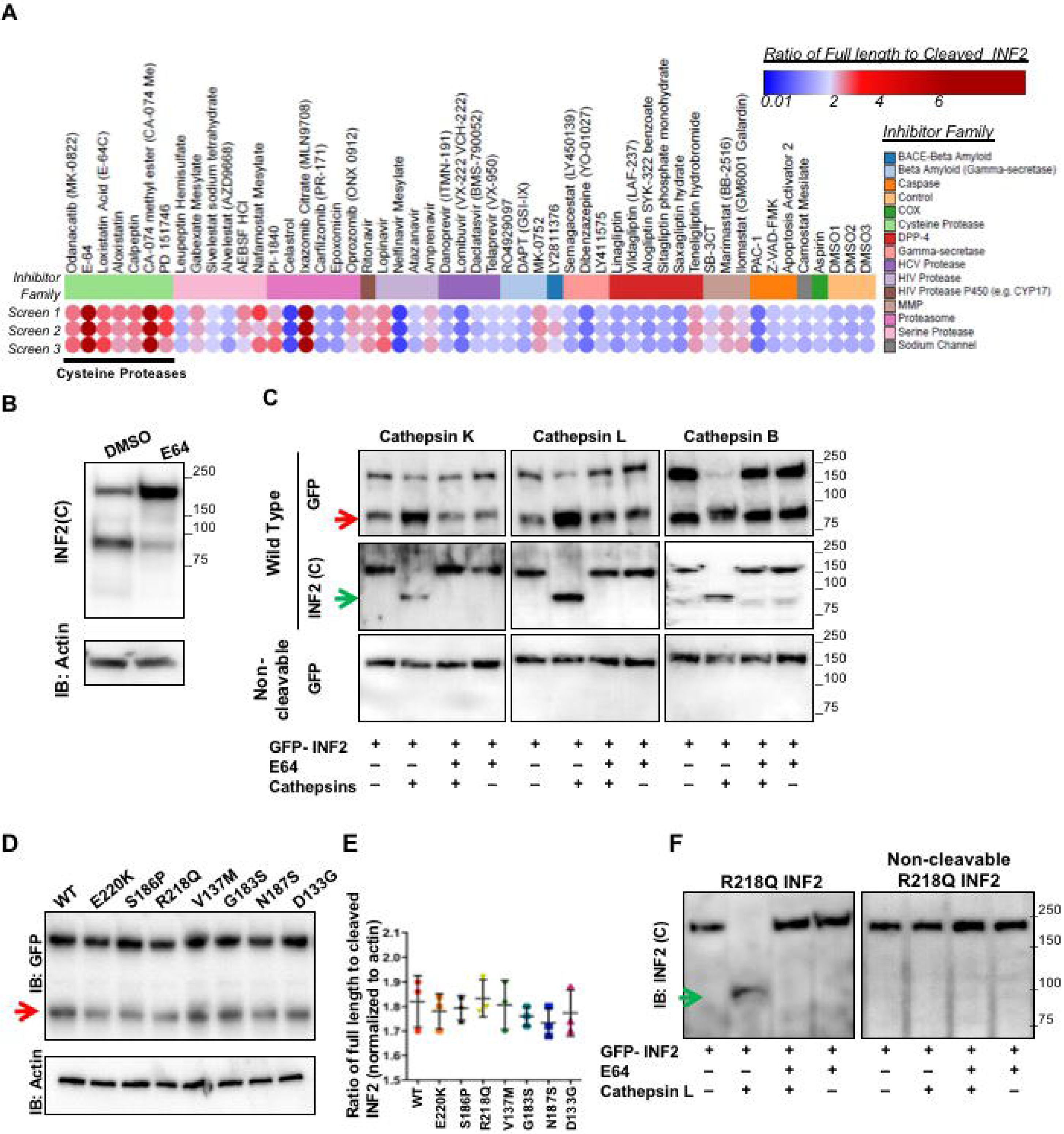
INF2-cleavage is mediated by cathepsins and is not affected by pathogenic mutations. (A) Screening of protease inhibitor library for effect on INF2 cleavage. Heat map shows the effects of different compounds on INF2 cleavage. Results from three independent sets of assays are shown. The cleavage levels with each treatment were calculated as ratios of full length to cleaved INF2 levels from INF2 immunoblots and plotted as a heat map, with color indicating inhibitor potency. Clustering of compounds into different protease inhibitor families showed that inhibitors of the cysteine protease family shared the ability to inhibit cleavage, most potently, E-64. (B) Representative immunoblot of INF2 cleavage inhibition by cysteine protease inhibitor, E-64. Green arrow indicates cleaved C-fragment. (C) *In vitro* cleavage assay. Immunoprecipitated GFP-INF2 or non-cleavable GFP-INF2 (Δ543-548) were incubated with the listed cathepsins, with or without E64. INF2 cleavage levels were then examined by immunoblot using N-terminal- (anti-GFP) or C-terminal-specific antibodies. All listed cathepsins cleaved GFP-INF2 but not the non-cleavable GFP-INF2 deletion mutant. The cleavage was inhibited by E-64. Red-arrow indicates cleaved N-terminal fragment. Green arrow indicates cleaved C-terminal fragment. Blots are representative of three independent experiments. (D) Immunoblot to detect cleavage of INF2 with different pathogenic mutations and other variants in the DID domain. (E) Quantitation of INF2 cleavage from three independent experiments is shown. No significant change in cleavage levels was noted in the presence of these INF2 mutations or variants in the DID domain (control vs. mutant and control vs. variant cleavages did not vary significantly (One-way ANOVA). (F) *In vitro* cleavage assay of INF2 with R218Q mutation. Immunoprecipitated GFP-R218Q INF2 or non-cleavable GFP-R218Q INF2 were tested for cleavage using cathepsin L. Immunoblot analysis showed E-64 inhibitable cleavage of GFP-R218Q INF2, but not GFP-R218Q INF2.

Next, to determine if INF2 cleavage is altered in the presence of FSGS-associated INF2 mutations, we assayed cleavage of full-length GFP-INF2 containing either FSGS-causing mutations (R218Q, E220K, S186P) or other variants predicted *in silico* to be either structurally permissive (V137M, G187S) or non-permissive (N183S, D133G), identified from the Exome Aggregation Consortium EXAC database. Transient overexpression of these constructs showed that comparable levels of cleavage occurs between control INF2 (wild-type) and all of the tested INF2 mutants and variants (Figure 3D and E). Similar results were observed using an in vitro cleavage assay with the R218Q mutant GFP-INF2 (Figure 3F), indicating that regulation of INF2 cleavage is not altered by the presence of pathogenic mutations. These results further suggest that the altered INF2 N-terminal antibody staining in human R218Q-diseased tissue sections cannot be attributed to altered proteolytic cleavage of INF2, but must be attributed to other mechanisms of altered INF2 N-terminal fragment localization. A similar manifestation of altered N-terminal fragment antibody staining in other disease conditions further indicate that mechanisms of altered localization are important in the disease (Supplementary Figure 2).

### INF2 N-fragment localization to the cell membrane is altered in the presence of disease-causing mutations

To evaluate INF2 fragment localization, we transiently overexpressed GFP-tagged INF2 N-terminal fragments (wild-type and R218Q), the INF2 C-terminal fragment, full-length INF2-CAAX (wild-type and R218Q), and non-cleavable INF2-CAAX (wild-type and R218Q) in mouse INF2 KO podocytes. The C-terminal fragment, the wild-type full-length INF2-CAAX, and the non-cleavable INF2-CAAX all showed prominent localization to endoplasmic reticulum (ER)-rich regions, while the wild-type N-fragment co-stained with cortactin at plasmalemmal regions (Figure 4A and B). A similar difference between N-fragment and C-fragment localization was observed when we stably overexpressed the HA-tagged N- or C- fragments in human podocytes, indicating that N-fragment localizes preferentially to plasma membrane regions in cells (Figure 4C and D). Importantly, we noted that the pathogenic FSGS mutation, R218Q, caused redistribution of the INF2 N-fragment from its normal plasma membrane-like locations to a diffuse pancellular staining pattern. Taken together, these results show that INF2 cleavage itself is not affected by the presence of disease-causing mutations, but that post-cleavage, localization of the N-fragment is dramatically altered with the pathogenic mutation.

**Figure 4:**
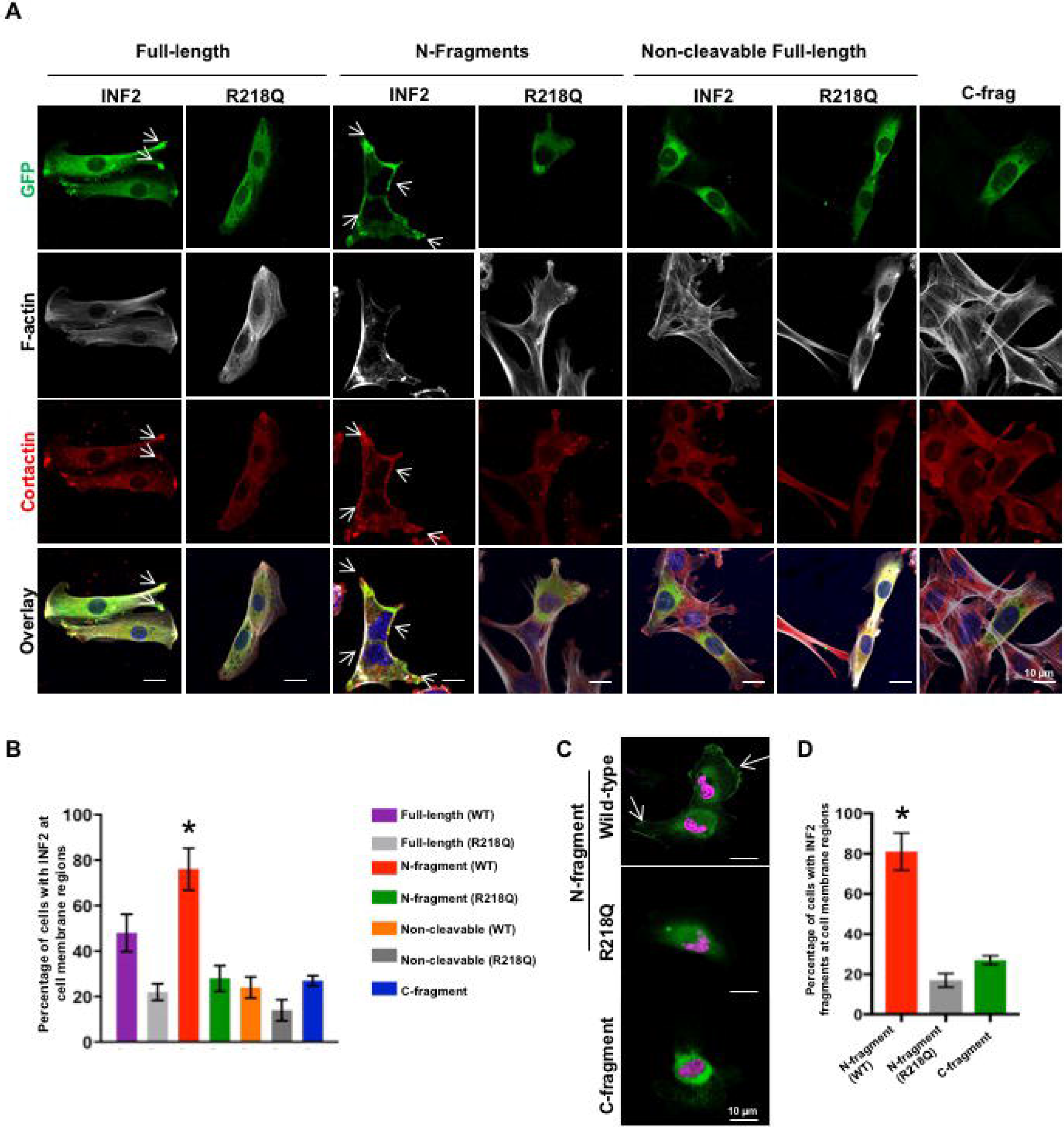
INF2 fragments exhibit differential localization in podocytes. (A) Localization of INF2 fragments in mouse podocytes. GFP tagged N-fragment, full-length, non-cleavable forms of wild-type or mutant INF2 (R218Q) and C-fragment of INF2 were transfected in INF2 KO podocytes and examined for their localization by co-staining with cortactin (red) and F-actin (grey). Wild-type N-fragment exhibited cell membrane colocalization with cortactin (white arrow highlights), while the C-fragment localizes to the cell body in an ER-like pattern. The R218Q mutation significantly altered the membrane localization of N-fragment leading to a diffuse cytoplasmic distribution, with loss of cortactin colocalization at the cell membrane. No membrane localization was observed for the non-cleavable forms of INF2, with or without R218Q. Scale 10 µm. (B) Quantitation of membrane localization of different INF2 expression forms (* p<0.01 GFP-N-fragment vs. all other groups; one-way ANOVA and Turkey’s multiple comparison test. n>100 for each group.). (C) Localization of INF2 fragments in human podocytes. HA-tagged INF2 N-fragment or C-fragment were expressed in human podocytes and localized; (Green, HA tag; Magenta, Nucleus). N-fragments expressed in human podocytes containing endogenous INF2 exhibited similar localization to membrane regions (white arrow highlights) along with some cell body staining, while the C-fragment localization was in an ER-like pattern. Membrane localization of N-fragment is altered by the R218Q mutation. Scale 10 µm. (D) Quantitation of cells with membrane-localized INF2 (*p<0.01, HA-N-fragment vs. HA-C-fragment or HA-R218Q fragment; one-way ANOVA and Turkey’s multiple comparison test. n>100 for each group.).

### INF2 N-fragment restores impaired cell spreading by antagonizing mDIA signaling

We next evaluated whether the cleavage-induced differential localization of the two proteolytic fragments of INF2 regulates any specific cell functions, and if INF2 N-terminal fragment function might be altered in the presence of INF2 mutations. Our previous studies have shown that both the mutant podocytes from R218Q knock-in mice and siRNA-silenced INF2 human podocytes exhibit impaired cell spreading (26, 30). These studies suggest that INF2 has a role in cell spreading that is altered in the presence of disease-causing mutations. We hypothesized that the cleaved N-fragment may normally help mediate the cell spreading function of INF2 and help maintain (or restore) normal podocyte structure.

To examine this, we first tested whether mouse INF2 KO podocytes and human INF2 CRISPR KO podocytes recapitulate the impaired cell spreading previously noted in other systems (Figure 5A). Consistent with previous observation, we found that both mouse INF2 KO podocytes and human INF2 CRISPR KO podocytes exhibited impaired cell spreading compared to their respective controls (Figure 5A). We next tested whether presence of the INF2 N-fragment restores normal cell spreading. As depicted in Figure 5B, we transfected mouse INF2 KO podocytes with different GFP-INF2 constructs to achieve equivalent expression levels and evaluated possible restoration of cell spreading. We found that expressing wild-type N-fragment in INF2 KO podocytes restored normal cell spreading in the INF2 KO cells. Normal cell spreading was partially restored by expression of the wild-type full-length INF2, but not by the non-cleavable form of full-length INF2. In contrast, none of the R218Q-mutant constructs produced any recovery in cell spreading. Similar effects on cell spreading were noted in human INF2 CRISPR KO podocytes transiently overexpressing wild-type INF2 N-fragment but not in podocytes overexpressing the R218Q mutant N-fragment (Figure 5C).

**Figure 5:**
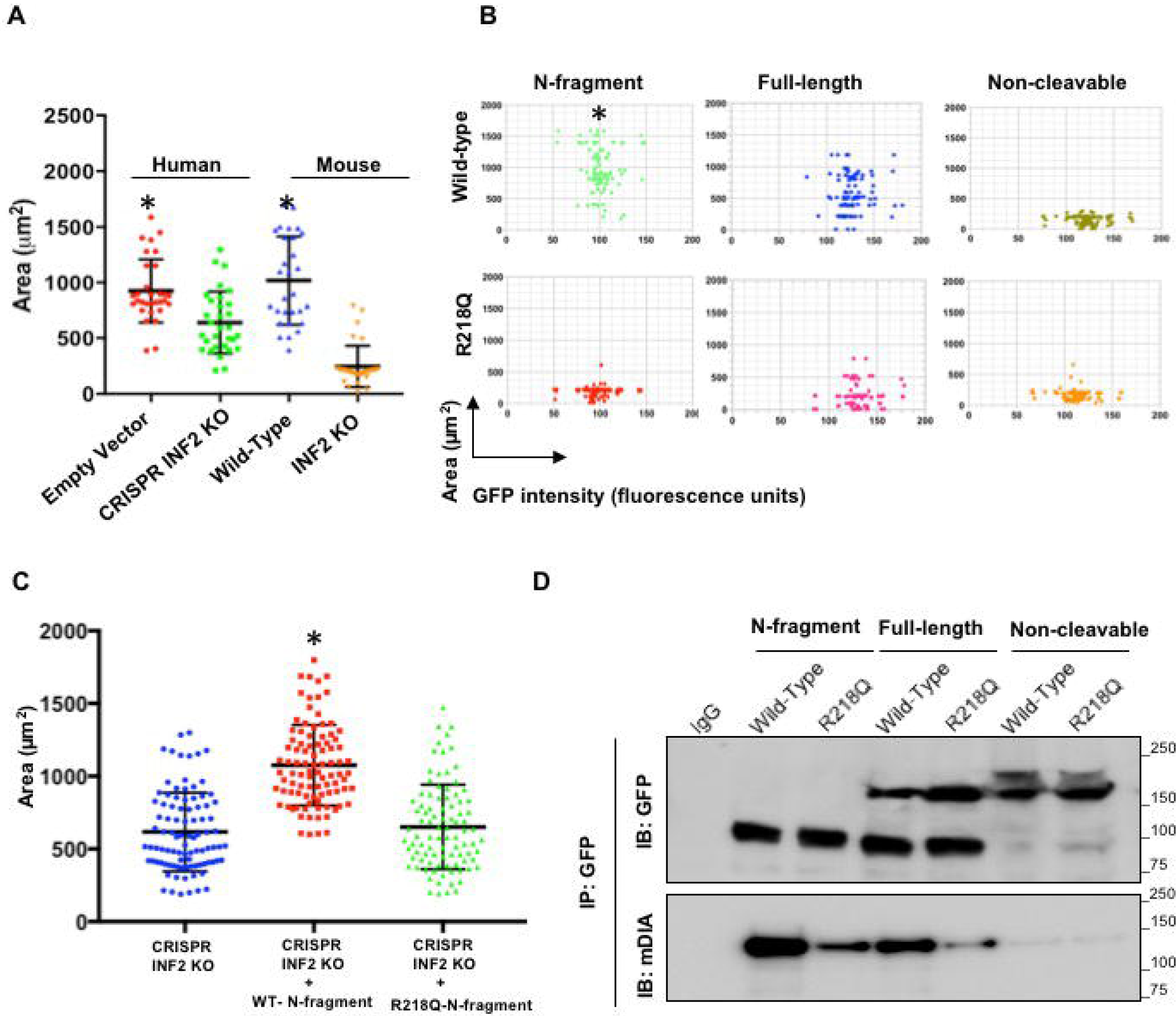
INF2 N-fragment restores cell spreading in association with mDIA interaction. (A-C) Cell spreading assays. (A) Mouse INF2 knockout podocytes and human INF2 CRISPR knockout podocytes were compared to their respective control cells for ability to spread upon attachment. The relative area covered after 45 minutes of cell spreading was calculated and analyzed for differences among groups (n> 100 cells for each group). Lack of INF2 in either mouse or human podocytes was associated with impaired cell spreading. (* p<0.001 EV vs. CR; WT vs. KO. One-way ANOVA and Turkey’s multiple comparison test. n>50 for each group). (B) Mouse INF2 knockout cells were transfected with GFP-tagged INF2 N-fragment, full-length INF2, or non-cleavable forms of wild-type or mutant INF2 (R218Q) and examined for restoration of normal cell spreading. GFP-tagged INF2 N-fragment expression significantly increased relative cell spreading area, whereas neither mutant nor non-cleavable forms of INF2 restored normal cell spreading (*p<0.01, N-fragment vs. other groups; one-way ANOVA and Turkey’s multiple comparison test. n>100 for each group). (C) Human INF2 CRISPR knock out podocytes were transfected with GFP-tagged wild-type or R218Q mutant N-fragment and spreading area was quantified. Wild-type N-fragment restored cell spreading whereas the R218Q mutant form did not. (* p<0.01, wild-type vs. R218Q; one-way ANOVA and Turkey’s multiple comparison test. n>100 for each group). (D) Co-immunoprecipitation of INF2 with mDIA. Cells transfected with GFP tagged INF2 N-fragment, full-length INF2, and non-cleavable forms of wild-type or mutant INF2 (R218Q) were subjected to GFP-pulldown and blotted for mDIA and GFP. GFP immunoblot confirmed successful immunoprecipitation of transfected INF2 constructs. mDIA immunoblot showed an interaction of mDIA with wild-type N-fragment and full-length INF2. Mutant forms of INF2 showed significantly decreased interaction, while non-cleavable forms of INF2 failed to interact with mDIA. Each immunoblot is representative of three independent experiments with similar results.

Our earlier studies showed that impaired podocyte cell spreading in the presence of INF2 mutations was associated with aberrantly increased mDIA signaling (26, 30), which we attributed to loss of an inhibitory interaction between INF2-DID and mDIA-DAD (19). Therefore, we examined whether the cleaved INF2 N-fragments interacted with mDIA in order to correlate our cleavage-dependent cell spreading observations with earlier findings suggesting a specific INF2-mDIA interaction {REFSXXX}. We observed that that wild-type INF2 N-fragment interacted with mDIA, and that this interaction was significantly reduced in the presence of R218Q mutant INF2. We observed a similar trend using full-length wild-type and R218Q mutant INF2 (Figure 5D). More importantly, the non-cleavable form of INF2 (both wild-type and R28Q mutant) did not interact with mDIA. These results indicate that INF2 cleavage mediates the antagonistic effect of INF2 DID on mDIA and that the INF2 N-fragment specifically promotes cell spreading. In the presence of an FSGS-associated INF2 point mutation, the ability of the N-terminal fragment to promote cell spreading is lost.

## DISCUSSION

Multiple studies have indicated that INF2, through its effects on actin and microtubules, can modulate mitochondrial fission, calcium uptake, vesicle trafficking, T-cell polarization, and placental implantation, among numerous other cell processes (23, 24, 31–34). The mechanisms by which FSGS-associated (and Charcot-Marie-Tooth associated) mutations cause disease is not well understood. In particular, explanations remain unclear for the restricted localization of these mutations to the INF2 N-terminal region, possible manifestations of differential evolutionary pressures experienced by the N and C terminal portions of INF2 gene products (8, 9, 16).

In this study, we have demonstrated that INF2 undergoes a proteolytic program that can cause the DID in the N-terminal region of INF2 to function independently of the DAD in the INF2 C-terminal region (Figure 6). As all the human *INF2* mutations identified to date have been confined to the DID-containing N-terminus, this observation suggests that N-fragment dependent functions must be integral for glomerular structure and function, and provides a potential clue to why all mutations are limited to the DID region. In addition, we observed that INF2-induced cell spreading and the interaction of mDIA with INF2 are cleavage-dependent properties and are both significantly reduced in the presence of a disease mutation. These results are consistent with our earlier findings on INF2 and mDIA-related cell spreading effects (26, 30). In this model, INF2 acts downstream of RhoA and inhibits mDIA-related actin cytoskeletal rearrangements. By doing so, INF2 promotes formation of lamellipodial structures and facilitates the trafficking of proteins and lipid raft structures to the cell surface. The present study suggests that this model is dependent on proteolytic cleavage of INF2. Consistent with the model that increased mDIA activity leads to disease, another recent study indicated that loss of mDIA1 can preserve glomerular function in a model of diabetic kidney disease (35).

**Figure 6:**
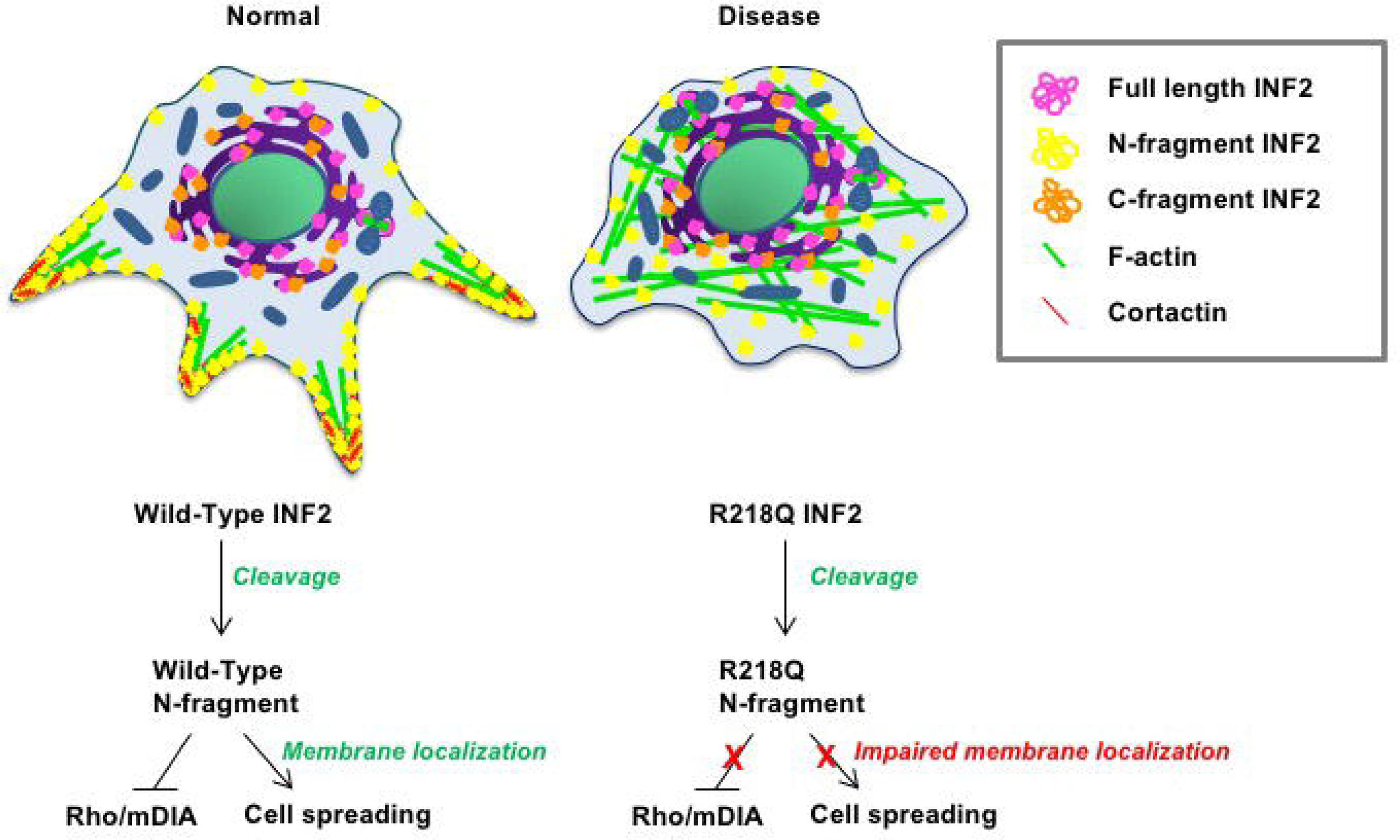
Summary Model: Cleavage-dependent unique functions of DID domain are altered due to mutations. INF2 is cleaved by cathepsins into two fragments, separating the DID and DAD regions. The cleaved DID-containing N-fragment localizes to cell membrane regions and functions to promote cell spreading, perhaps by counteracting mDIA signaling. However, with the R218Q mutation, both cell spreading and mDIA interaction are impaired. In one possible model of INF2-mediated disease, insults that cause podocytes to lose slit diaphragm integrity may also activate cathepsins, promoting INF2 cleavage and allowing the N-terminal DID-containing region of INF2 to help restore podocyte structural integrity. With pathogenic mutations, loss of proper localization and lack of DID function may lead to persistent injury.

Our analysis of *INF2* transcript variants indicates that podocytes express an additional transcript that includes just the 5’ portion of the gene, encoding only the DID. However, we were unable to detect a specific immunoreactive protein product of this transcript in podocyte lysates by immunoblot. This short isoform may function as a regulatory RNA, or may be translated and/or accumulate only in specific conditions yet to be identified. Additional analysis of this short isoform will be needed to identify its possible functional roles.

Prior *in vitro* studies of full-length INF2 have indicated that INF2 is normally kept in an auto-inhibited state and is activated under spatial and temporal control (15, 23). Our studies show that INF2 cleavage physically separates the N and C terminal portions of the polypeptide and activates at least some DID-mediated functions of INF2.

Tamura et al. reported, using a C-terminal antibody-based immunohistochemical analysis, that INF2 in normal glomeruli is present in the podocyte cell body and major processes, with decreased or absent expression in these sites in FSGS (36). We have observed localization of INF2 C-terminal antibody staining in normal glomeruli. However, with the identification of INF2 cleavage, we suggest that caution is required in correlating INF2 staining intensities with disease states. The amount of INF2-cleavage and the stability of the two fragments may vary in different disease states. Nevertheless, the results presented here point to a specific role for the N-terminal region in podocyte foot processes. This activity may be lost in FSGS caused by N-terminal INF2 mutations.

All FSGS-associated mutations (with or without associated CMT) localize to the INF2 N-terminal region (37). This suggests that the N-terminal region is important for the integrity of the foot process and slit diaphragm structures, and suggests that INF2 cleavage has *in vivo* significance, particularly in the context of disease. We note that the loss in N-fragment immunostaining intensity in the INF2 R218Q mutant kidney is not limited to foot process structures. A grossly reduced staining intensity of this fragment is also noted in the cell body. By contrast, the staining intensity of the INF2 C-fragment remains unaltered. Although speculative, this could result from: (1) increased cleavage leading to increased fragment formation, and/or (2) presence of a mutation affecting the stability of mutant N-fragment but not the C-fragment, thus causing a loss of total N-fragment intensity but not C-fragment intensity. Future studies designed to determine changes in INF2 cleavage levels in the course of the disease and further analysis of the stability of normal and mutant forms of INF2 (full length and fragment) should help clarify these questions.

Our experiments have shown that cathepsin proteases are primarily responsible for INF2 cleavage. A diverse set of stimuli, including transient injury events, can upregulate and activate cathepsins in the kidney (38). Multiple studies have indicated that cathepsin proteases can inactivate CD2AP and other proteins essential for podocyte structure and function (39). Proteolytic processing of α-actinin-4, podocin, and other proteins by cathepsins has recently been suggested to facilitate maintenance of podocyte structure and filtration barrier integrity (40). It is thus interesting to note that INF2 is also a target of cathepsin proteases. Studies of human samples indicate evidence of INF2 cleavage in normal kidney (Figure 1). Previous studies in mouse models indicate that INF2 functions as an injury response protein, facilitating podocyte recovery (30). Given that cathepsins are upregulated in transient injury conditions, it is likely that cathepsin-mediated INF2 cleavage regulates N-terminal fragment activity. Thus, taken together, it is intriguing to propose that upon glomerular injury, increased cathepsin activity causes increased proteolytic cleavage of INF2, and that increased N-fragment activity might facilitate podocyte recovery (30). This hypothesized role of INF2 in podocyte repair would be lost in the presence of FSGS-causing mutations. Further studies are needed to evaluate this model.

Multiple *in vitro* and *in vivo* studies have shown that cell spreading and related cytoskeleton defects (e.g., loss of cortactin) are hallmarks of impaired INF2 function in cells (26, 30). Our study provides evidence that cleavage-generated INF2 N-fragment exhibits differential localization to membrane regions and regulates cell spreading functions, while C-fragment remains with an ER-like distribution throughout the cell body. Regulation of actin dynamics and cell spreading functions by formin family members is important for proper adherens junction assembly (41). Given that slit diaphragm structures of podocytes share morphological features similar to those of the adherens junction, the INF2 N-fragment may mediate or facilitate the actin dynamics required for proper slit diaphragm structure and function (42). Although, INF2 N-fragment mediated cell spreading defects provide one possible mechanism for linking mutant INF2 with FSGS, changes in the full gamut of INF2 activities will need exploration.

In summary, we have shown that INF2 undergoes proteolytic cleavage that can lead the N-terminal DID region to function independently of the C-terminal DAD region (Figure 6). In addition, we have observed cell spreading activity and mDia interaction that are cleavage-dependent and are significantly affected with by pathogenic mutation of INF2. These data provide a plausible explanation for the altered podocyte localization of INF2 N-fragments with FSGS-associated mutations. Future studies to define the full range of INF2 N-fragment and C-fragment functions in maintenance of glomerular podocyte structure and function will increase our understanding of the physiological roles of both INF2 cleavage and the resultant INF2 proteolytic fragments.in health and disease.

## MATERIALS AND METHODS

### Cell culture

Mouse podocytes generated from wild-type and INF2 KO C57BL/6 mice (Supplementary Figure 3) and human podocytes (43) were maintained in RPMI 1640 (ThermoFisher Scientific, Waltham MA) supplemented with 10% fetal bovine serum (ThermoFisher), Insulin-Transferrin-Selenium (ITS-100X) and 1% pen-strep (ThermoFisher). Human primary podocytes from Celprogen Inc. (Torrance, CA) were maintained per manufacturer’s instructions. Human embryonic kidney epithelial cells (HEK293T cells) were maintained in Dulbecco’s Modified Eagle Medium supplemented with 10% fetal bovine serum (ThermoFisher) and 1% pen-strep (ThermoFisher).

### Plasmid constructs and INF2 expression

For overexpression experiments, human INF2-CAAX isoform sequence corresponding to amino acids 1-547 (N-fragment), 548-1249 (C-fragment), and 1-1249 (full-length) were cloned into an EGFPC1 plasmid (N-terminal GFP tag) (Takara Bio, Mountain View, CA). For cleavage site validation experiments, FLAG-tag was inserted between the CAAX motif (CVIQ) and stop codon in the full-length construct (Cat No: 200523, QuickChange II Site-Directed Mutagenesis kit, Agilent Technologies, Lexington, MA). For removal of the INF2 cleavage site, amino acids 543-548 of INF2 were deleted in full-length CAAX constructs. Pathogenic mutations and variations in the INF2 sequence were introduced by a PCR-based mutagenesis (Cat No: 200523, QuickChange II Site-Directed Mutagenesis kit, Agilent Technologies). To achieve stable expression of fragments in human podocytes, N- and C-fragment were cloned into a lentiviral expression plasmid (Cat. No: PS100101, OriGene, Rockville, MD). For generating CRIPSR KO podocytes, gRNA (sense: TGCGCGCCGTCATGAACTCG; antisense: CGAGTTCATGACGGCGCGCA) targeting the DID domain region of INF2 was expressed using a lentiviral plasmid (Cat. no:5296, lentiCRISPRV2, Addgene, Boston, MA). All cloned plasmids were confirmed for the correct sequence by DNA sequencing (Genewiz, Boston, MA). For variations and mutant cleavage analysis, variations in the DID domain of INF2 were first extracted from Exome Aggregation Consortium (ExAC) and analyzed using a software tool, PROVEAN (Protein Variation Effect Analyzer), to group the variations as either structurally permissive and non-permissive variants. Representative variants for structurally permissive and non-permissive groups and pathogenic mutants were then cloned into EGFPC1 plasmid and overexpressed in cells.

### Co-immunoprecipitation

HEK293T cells were transiently transfected with the indicated INF2 constructs using the Lipofectamine 2000 (Invitrogen, Carlsbad, CA). After incubation for 24 h, the cells were lysed in 1% NP-40 lysis buffer (1% NP-40, 50mM Tris-HCl, 150mM NaCl, 5mM EDTA, pH 7.4) supplemented with protease inhibitors. Cell lysates were then incubated with mouse anti-GFP-tag antibody (MA5-15256, ThermoFisher) for 1 hr followed by 40 μL of protein A/G-magnetic beads for another 1 hr at 4°C. The co-immunoprecipitation of DIAPH1 (A300-078A, Bethyl labs, Montgomery, TX) was then analyzed.

### Immunoblotting

Cells or glomerular preparations were lysed in a RIPA buffer (Boston BioProducts, Ashland, MA) supplemented with a cocktail of protease and phosphatase inhibitors (Roche, Pleasanton, CA) and clarified by centrifugation at ~13000 g for 15 minutes at 4 ºC. Equal protein loads were separated on a 4-20% reducing gel, transferred to a Polyvinylidene difluoride (PVDF) membrane (Bio-Rad, USA), and probed with respective primary and secondary antibodies as follows: INF2 (N) (1:500) (A303-427A, Bethyl Labs); INF2 (C) (1:500) (20466-1-AP, Proteintech, Rosemont, IL); Podocin (1:1000) (P0372, Sigma, St Louis, MO); anti-rabbit (1:4000) (Santa Cruz Biotechnology, Dallas, TX). The membranes were then developed using a chemiluminescent-based substrate (Super Signal West Dura, ThermoFisher). Total beta-actin (1: 4000) (sc47778; Santa Cruz) level was used as a loading control.

### Chemical screen

To assess changes in INF2 cleavage with different protease inhibitors, human podocytes were treated with small molecules from a protease inhibitor library (SelleckChem LLC, Houston, TX). Cells were treated with a concentration of 10 μM for 24 hrs. Post incubation, cells were lysed in RIPA buffer and lysates examined for cleavage inhibition by INF2 immunoblot. The ratio of full-length INF2 to the cleaved C-terminal fragment was used to quantitate INF2 cleavage and inhibition of cleavage by the library compounds.

### INF2 isoform analysis

Total RNA of human podocytes was extracted per manufacturer’s instructions (Qiagen, Germantown, MD) and 1 μg RNA was used for cDNA synthesis (Transcriptor first strand cDNA synthesis kit, Roche). INF2 was then PCR amplified from cDNA using isoform-specific primers (Supplementary Figure 1) (Accuprime DNA Polymerase system, ThermoFisher). The PCR amplified product was gel-purified and examined for INF2-isoform-specific sequences using DNA sequencing (Genewiz).

### Cleavage site mapping and in silico INF2 cleavage analysis

HEK 293T cells were transiently transfected with the GFP-INF2-FLAG constructs using Lipofectamine 2000 (Invitrogen). After incubation for 24 hr, the cells were lysed in RIPA buffer and the clarified supernatant was subjected to immunoprecipitation by anti-flag M2 beads (Sigma) for 2 hr, followed by washing and elution by anti-flag peptide (Sigma). The eluates were run on a 4-20% denaturing gel, transferred to a PVDF membrane, and stained using Coomassie blue. The C-fragment band was cut, digested per vendor guidelines (Alphalyse, Palo Alto, CA) and subjected to N-terminal sequencing. To assess the cleavage sites in INF2 *in silico*, we used the PROSPER bioinformatics tool.

### In vitro cleavage assay

The *in vitro* cleavage assay was performed as described previously (39). Briefly, immunoprecipitated GFP-INF2 was diluted in a buffer containing 200 mM NaCl, 10 mM HEPES (pH 7.0), 2 mM EGTA, 1 mM MgCl2, and 1 mM DTT. When indicated, 100 μM E64 inhibitor (SelleckChem) was added. The reaction was initiated by addition of purified Cat L (0.1 μL), B (0.1 μL), or K (0.05 μL) enzyme (Sigma), and samples were placed at 37°C in a water bath for 15 minutes. Total assay volume was 25 μL. The reaction was terminated with the addition of 4x sample buffer (BioRad, Hercules, CA)

### Cell spreading assays

Cells were serum starved for two consecutive days at 0.5% FBS in RPMI1640 medium, then trypsinized and seeded in equal number on fibronectin-coated cover slips for 45 minutes. Cells were washed with ice-cold PBS buffer and processed for either fixation or lysis. The cell spreading area was measured after immunostaining fixed cells with anti-F-actin.

### Immunofluorescence

Cells were stained and imaged using confocal microscopy as described previously (30). Briefly, cells were fixed with 4% paraformaldehyde for 15 min, PBS-rinsed, and permeabilized 15 min with 0.5% Triton x-100. The fixed cells were then blocked with 5% BSA followed by sequential incubation with primary and secondary antibodies in blocking buffer. Following antibody incubations, cell nuclei were counterstained with Hoechst (dsDNA) (Invitrogen), mounted using ProLong Diamond Antifade (Invitrogen), and imaged by confocal microscopy (Zeiss LSM 510). All images were collected using ZEN lite 2.3 (black edition) and processed using ZEN lite 2.3 (blue version).

### Structured Illumination Microscopy (SIM) and Image analysis

SIM analysis of kidney biopsy section samples was performed using a Zeiss Elyra SP.1 system, as described previously (44). Briefly, z-stacks were acquired over a volume of 75.44 × 75.44 × 4 μm^3^ (length × width × depth) with a slice-to-slice distance of 0.13 micron. The 34 μm period grating was shifted five times and rotated five times on every frame. SIM processing post-image acquisition was performed in 3D stacks using Zeiss ZEN software with default processing parameter settings.

The SIM-processed frames were then converted to a maximum intensity projection image using ZEN software or individually analyzed for foot process organization and INF2 localization using FIJI software. The foot processes were tracked in z-stack frames by synaptopodin staining. Foot process regions at points orthogonal to the imaging frame were defined as “Regions Of Interest” (ROI) and used for analysis. The profiles of synaptopodin and INF2 fluorescence intensity were then plotted and examined for co-localization of peaks.

### INF2 knockout mouse

A neomycin cassette was introduced into the mouse *Inf2* gene locus by homologous recombination to generate an *Inf2* knockout allele in C57Bl/6 embryonic stem cells. The cassette targets the *Inf2* gene locus spanning the start site of *Inf2* transcription, causing a complete loss of INF2 expression. Correctly targeted embryonic stem cells were injected into 8-cell Swiss Webster mouse embryos. Embryos were cultured overnight and transferred into pseudo pregnant female mice 2.5 days post-coitus. F0 male mice were bred with C57BL/6 mice to generate KO mice (See Supplementary Figure 3). All mice were used for experiments after breeding for at least five generations.

### Statistical analysis

All statistical analyses were performed using one-way analysis of variance between test-groups. When statistical significance was seen, Turkey’s multiple comparison test was used to find group differences. Statistical significance was set at a minimal value of p <0.05. All calculations were made using GraphPad Prism Version 7, and all values were reported as means ± standard deviation.

### Human kidney studies

Human kidney biopsy material was obtained from Beth Israel Deaconess Medical Center (BIDMC) or outside institutions in accordance with a protocol approved by the Institutional Review Board at BIDMC.

## AUTHOR CONTRIBUTIONS

Balajikarthick Subramanian: Conceived the idea, managed the study, performed the experiments, analyzed the data and wrote the manuscript. Justin Chun: Performed the experiments, analyzed the data and wrote the manuscript. Chandra Perez and Paul Yan: Managed the mouse line, Performed the experiments: Isaac Stillman: Analyzed the human tissue data. Henry Higgs, Seth L. Alper and Johannes Schlondorff: Analyzed the data and edited the manuscript. Martin R. Pollak: Conceived the idea, analyzed the data and wrote the manuscript.

## ACKNOWLEDGEMENTS

We thank the Harvard Center for Biological Imaging, Harvard University, Cambridge, MA for help with super-resolution imaging and analysis. We thank BIDMC confocal core facility, Boston, MA for help with immunofluorescence imaging and analysis. This work was supported by NIH grant RO1DK088826. Balajikarthick Subramanian was supported in part by NIH T32 award DK007199.

---

**Supplementary Figure 1.**
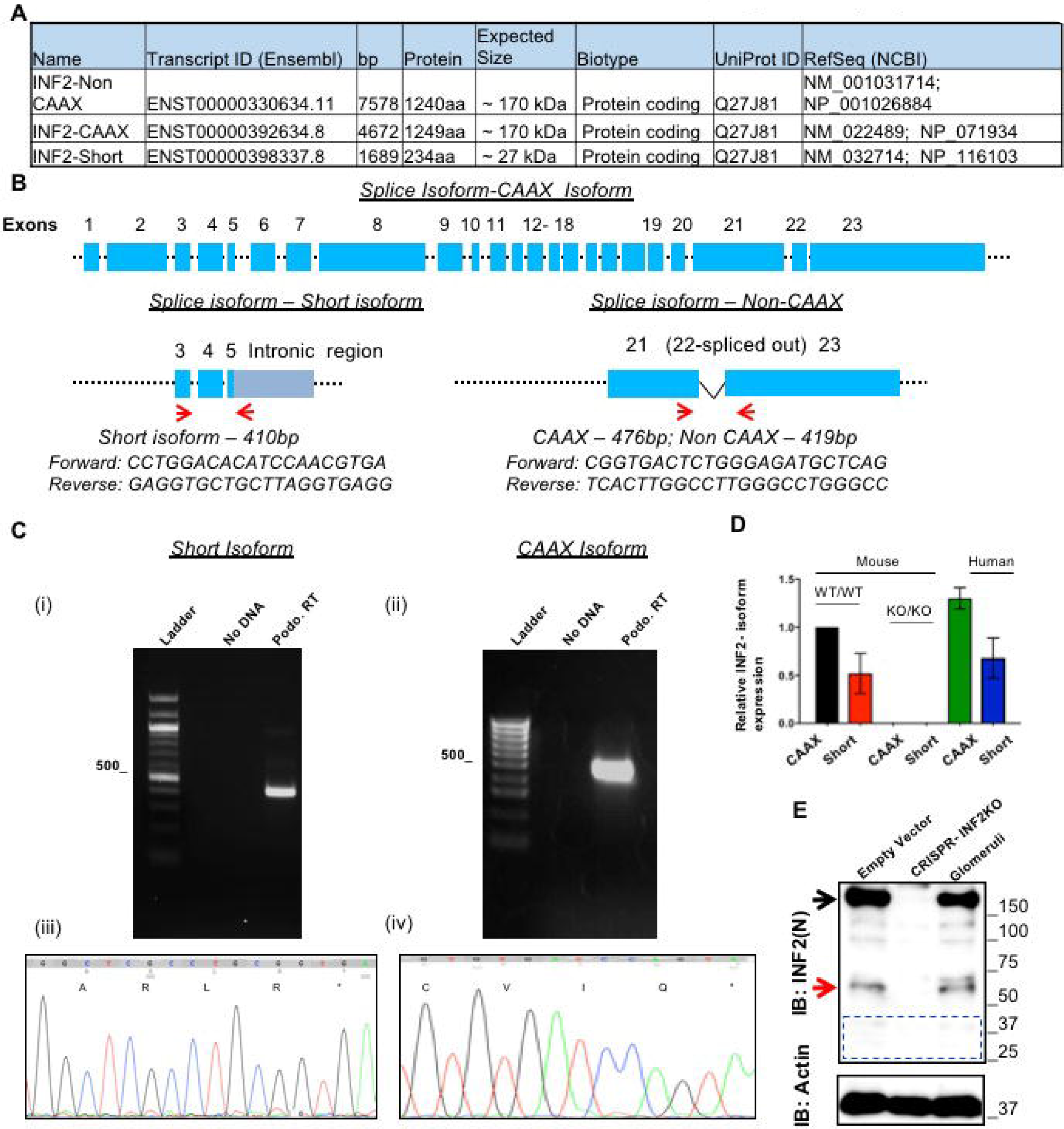
Splice isoform analysis of INF2 in podocytes. (**A**) The protein-coding isoforms of INF2 from ENSEMBL and NCBI databases that span the DID domain region. (**B**) Schematic of INF2 isoforms with exons (not drawn to scale) and primer binding sites (red arrows). (**C**) Isoform expression analysis in immortalized human podocytes. DNA gel electrophoresis analysis confirms single PCR product amplification of expected size from each transcript (short isoform: 410 bp; INF2-CAAX: 476 bp). Sequencing of PCR-amplified products confirms isoform subtypes, as shown in sequence traces with amino acid translations (iii) Short-isoform (* stop codon); (iv) INF2-CAAX (CAAX motif (CVIQ) is shown). (**D**) Confirmation of similar isoform expression patterns in mouse and human glomeruli. Mouse: WT/WT (Wild-type), KO/KO (Inf2 knock out). Expression data are normalized to nephrin expression and reported relative to mouse CAAX isoform. (**E**) Polypeptide isoform analysis by Immunoblot. Total cell lysates from human immortalized podocytes (empty vector and INF2 CRISPR KO) and human glomeruli were probed using INF2 N-terminal antibody. Black arrow: full-length INF2; red arrow: cleaved N-fragment); blue highlight: expected region for short isoform). No immunospecific band was reproducibly seen in the blue highlighted region.

**Supplementary Figure 2.**
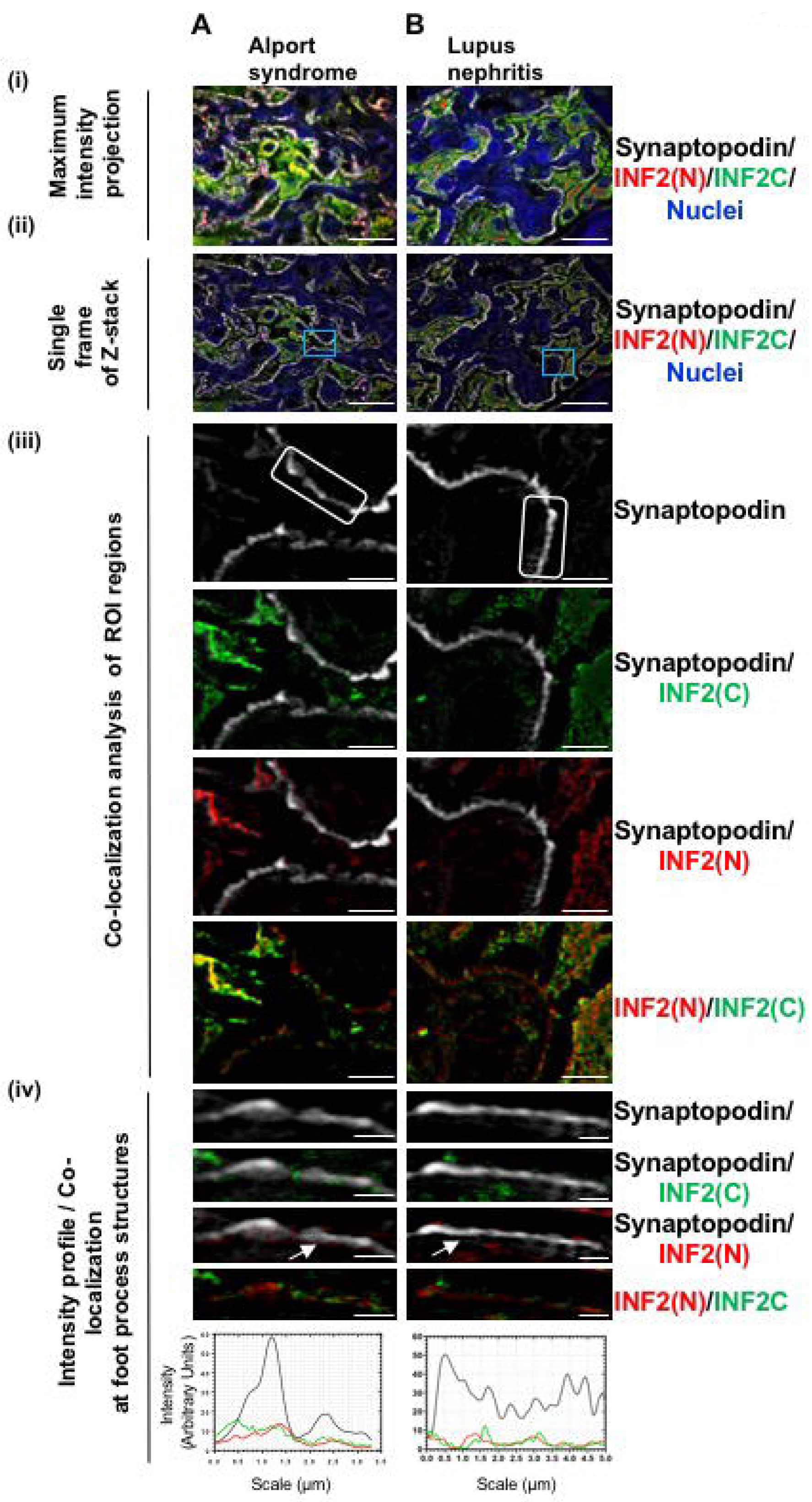
Altered localization of INF2 N-fragment in diseased glomeruli. Kidney-biopsy sections from individuals with (**A**) Alport Syndrome and (**B**) Systemic lupus erythematosus were stained for synaptopodin (grey), INF2 N-terminus (N, red), INF2 C-terminus (C, green), and nuclei (blue). Representative glomerular structures are shown. (**i & ii**) SIM processed low magnification micrographs. Scale bar 20 μm. (**i**) Maximum intensity projection of 3D Z-stack optical frames. (**ii**) A representative single optical frame of the 3D Z-stack shown in (i). Blue box highlight: region of interest (ROI) used for co-localization analysis. (**iii**) Co-localization analysis of INF2 with synaptopodin. White box highlights foot process structure. Scale bar 2.5 μm. (**iv**) Co-localization analysis and corresponding intensity profiles of INF2 and synaptopodin in foot process structures. Scale bar 0.7 μm. Similar to the R218Q INF2-mutant patient, INF2-N terminus co-localization is significantly reduced in effaced foot process structures (arrow highlights in Aiv-Biv). The intensity profile confirms lack of co-localized fluorescent signal intensity.

**Supplementary Figure 3:**
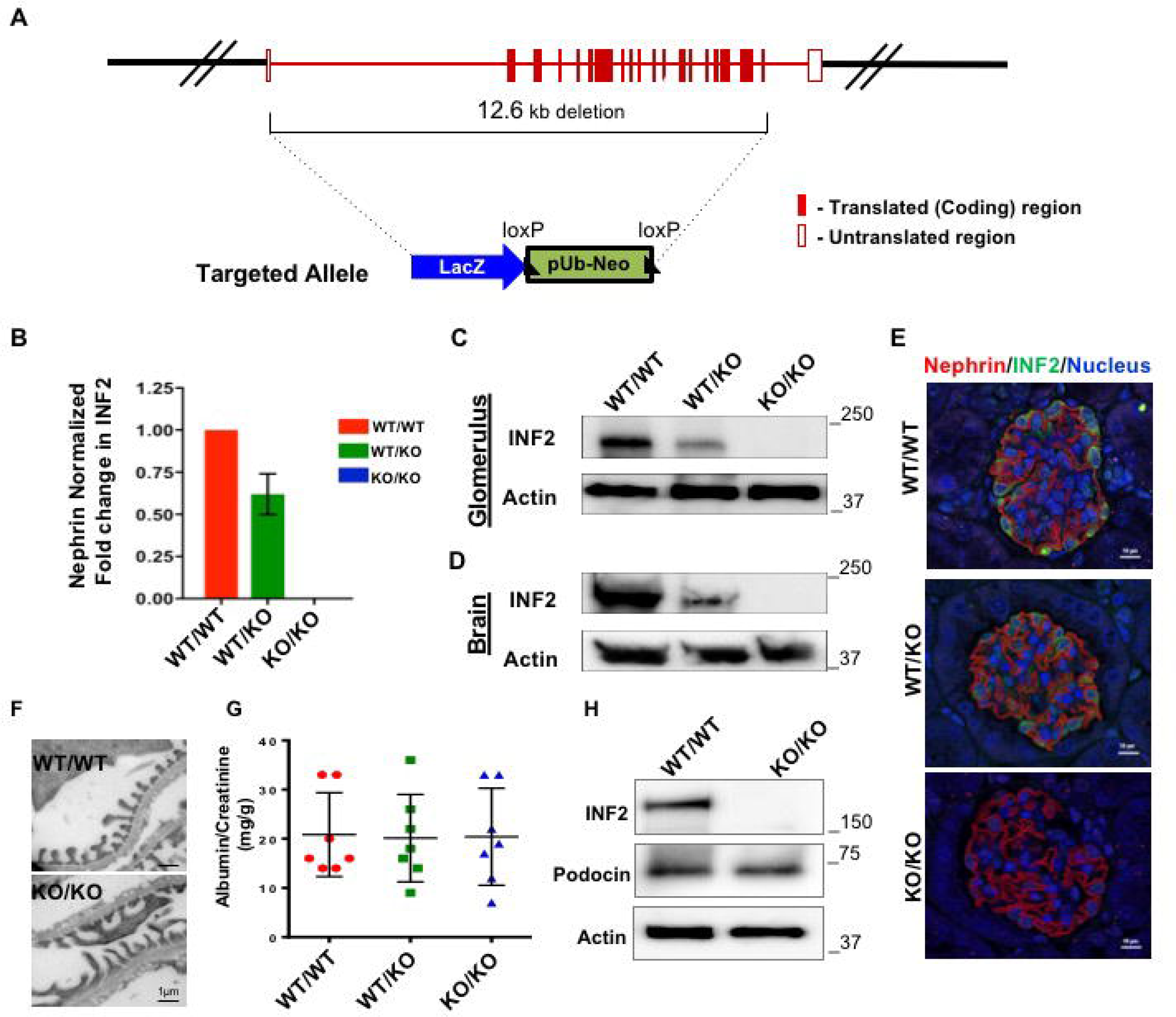
Generation and characterization of *Inf2* knockout mouse and podocytes. (**A**) Schematic of *Inf2* knockout mouse targeting strategy. The neomycin cassette was introduced into the mouse *Inf2* gene locus by homologous recombination to generate an *Inf2* knockout allele in C57Bl/6 embryonic stem cells. The cassette targets the *Inf2* gene locus spanning the *Inf2* transcriptional start site, causing complete loss of *Inf2* expression. (**B-E**) Expression analysis of *INF2* in glomeruli. (**B**) Real-time PCR analysis of *INF2* from RNA extracted from glomerular preparations of wild-type (WT), heterozygous (WT/KO), and homozygous (KO/KO) mice. Expression data is normalized to nephrin transcript levels. Results are from three independent glomerular preparations. (**C&D**) Immunoblot of INF2 from WT, WT/KO, and KO/KO glomeruli (**C**) and brain (**D**), each representative of three independent experiments. (**E**) Immunofluorescence staining of INF2 in WT/WT, WT/KO, and KO/KO mouse glomeruli. Complete loss of INF2 mRNA and protein was noted in the homozygous knockout (KO/KO) samples, while dose-dependently reduced expression was noted in the heterozygous knockout (WT/KO). (**F**) Electron microscopy analysis of *Inf2* KO mouse. Comparison of electron micrographs from both WT and KO mouse revealed no significant change in ultrastructural organization of podocyte foot processes in the *Inf2* KO mouse. (**G**) Comparison of urine albumin/creatinine ratios (ACR) from Inf2 WT, WT/KO, and KO/KO mice revealed no significant increase in ACR associated with either hetero- or homozygous KO allele. (**H**) Characterization of podocyte preparation from *Inf2* knock out mouse. Podocytes outgrown from glomeruli of WT/WT and KO/KO mice were immortalized using temperature sensitive large T-antigen. Differentiation of podocytes was induced at 37 C for 10 days. The presence of KO allele and the podocyte origin of cells were respectively confirmed by INF2 and podocin immunoblots with actin as loading control. Podocin expression was observed in both cell types, while INF2 expression was limited to the wild-type group. The immunoblot shown is representative of three independent experiments.

## Notes

**DISCLOSURE OF CONFLICTS OF INTEREST**: The authors have declared that no conflict of interest exists

